# How sequence context-dependent mutability drives mutation rate variation in the genome

**DOI:** 10.1101/2021.07.20.453104

**Authors:** Madeleine Oman, Aqsa Alam, Rob W. Ness

**Affiliations:** Dept of Ecology and Evolutionary Biology, University of Toronto, Toronto, Canada; Dept of Biology, University of Toronto, Mississauga, Canada; Dept of Cell and Systems Biology, University of Toronto, Toronto, Canada

## Abstract

The rate of mutations varies >100-fold across the genome, altering the rate of evolution, and susceptibility to genetic diseases. The strongest predictor of mutation rate is the sequence itself, varying 75-fold between trinucleotides. The fact that DNA sequence drives its own mutation rate raises a simple but important prediction; highly mutable sequences will mutate more frequently and eliminate themselves in favour of sequences with lower mutability, leading to a lower equilibrium mutation rate. However, purifying selection constrains changes in mutable sequences, causing higher rates of mutation. We conduct a simulation using real human mutation data to test if (1) DNA evolves to a low equilibrium mutation rate and (2) purifying selection causes a higher equilibrium mutation rate in the genome’s most important regions. We explore how this simple process affects sequence evolution in the genome, and discuss the implications for modelling evolution and susceptibility to DNA damage.

## Introduction

Mutation introduces the genetic variation that facilitates adaptive evolution but can also give rise to diseases such as cancer and age-related illnesses (Alexandrov et al. 2013; Bae et al. 2018). Although mutation is random, the relative rate of mutation at each position, or mutability, is not uniform across the genome. Variation in mutability has been demonstrated at multiple scales and the rate at which mutations occur can vary from site to site across the genome more than 100-fold (mammalian genome mutation rate variation reviewed in Ellegren et al. 2003). Thus, individual sites and regions across the genome have the potential to participate in evolution and genetic pathologies at varying rates. Analysing the drivers and patterns of mutability variation can help us understand the mutagenic mechanisms of human disease and the forces that generate the variation available to evolution.

Investigating how molecular processes and genomic properties predict mutation is an active area of study (Chen et al. 2017; Supek and Lehner 2019), and evidence suggests that the strongest predictor of mutation rate is the sequence itself (Michaelson et al. 2012; Sung et al. 2015). Although the mechanisms are not fully understood (Sung et al. 2015), we know that a given genomic position’s mutability is strongly influenced by not only the base itself but also the sequence at adjacent sites (Blake et al. 1992; Hess et al. 1994; Aggarwala and Voight 2016). Studies examining variation in the mutation rate among the 64 possible trinucleotide sequences have shown that the most mutable trinucleotides mutate at rates up to 75-fold higher than the least mutable (Sung et al. 2015). Therefore, if mutation rate variation among sites is influenced by sequence context, the uneven rate of mutation across the sequence should lead to changes in the underlying mutation rate variation over long periods of time.

The effect of mutation events on local mutability raises interesting questions about how sequence context and mutation rate might interact over long periods. We hypothesize that if high mutability sequences mutate more often, they are likely to persist in the genome for less time than lower mutability sequences that are more stable. Over time this bias would lead to an enrichment of lower mutability sequences and an overall depression of mutation rate. However in conserved sequences, we predict that purifying selection would prevent the fixation of mutations that compromise functionality. By restricting the sequence’s evolution to a lower mutability state, purifying selection may indirectly maintain some sites with higher mutability. Thus, we expect constrained sequences may evolve higher mutability than neutral regions.

In this study we conduct a long-term simulation of DNA sequence evolution under a trinucleotide-based mutation model generated from empirical human germline mutation data. We study the influence of sequence context mutability variation on mutation rate evolution. We test our two main predictions that; (1) DNA will evolve to a low equilibrium mutation rate and (2) purifying selection will constrain the change of sequence resulting in higher equilibrium mutation rates in the genome’s most important regions. We also explore how this change in sequence composition, driven by mutability variation, manifests in the frequencies of codons in protein coding genes under purifying selection.

## Results & Discussion

### The mutability model

To simulate how sequence context influences the evolution of mutation rate variation, we generated a model of trinucleotide mutability based on ~108k *de novo* mutations (Jónsson et al. 2017). Each trinucleotide in our model contains the mutability, as well as the probabilitys it mutates into each of the three potential descendant trinucleotides. Our model captured a large amount of mutability variation among the 32 trinucleotide sequences, with a 27-fold difference between least (G<u>A</u>A) and most (A<u>C</u>G) mutable trinucleotide. As expected, our mutability model shows that CpG trinucleotides have on average 10-fold higher mutability than non-CpG trinucleotides (Hwang and Green 2004; Hodgkinson and Eyre-Walker 2011). It is well documented that CpG sites have the highest mutation rate in humans and many other species, primarily due to deamination mutations at methylated cytosines (Coulondre et al. 1978; Razin and Riggs 1980). However, non-CpG bearing trinucleotides still show considerable variation, with a 3.7-fold difference between the lowest (G<u>A</u>A) and highest (G<u>G</u>T) mutability trinucleotides. Even though our model is based on human mutations, similar variation is seen in other species. In addition to the 75-fold trinucleotide mutability variation found in mismatch-repair deficient *B. subtilis* (Sung et al. 2015), we calculated similar trinucleotide mutability models for other organisms with sufficiently large mutation accumulation (MA) datasets and found 20-fold global mutability variation in *C. reinhardtii* (Ness et al. 2015, n=6843), 12-fold in *S. cerevisiae* (Zhu et al. 2014, n=867), and 55-fold in *M. musculus* (Lindsay et al. 2019, n=764). Therefore, although the results we present here are focused on a human model, it seems reasonable that similar processes could apply across a broad range of organisms.

### Does mutability of neutral sequence reduce over time?

Our first prediction was that sequence will evolve to a lower mutability state at equilibrium because high mutability trinucleotides will mutate more often, and over time lower mutability trinucleotides will become more common. To test our prediction, we simulated the evolution of a neutral (non-coding) sequence under our mutability model. In support of our prediction, starting from random sequence, the average mutability decreased by 53% at equilibrium. We would caution that the exact magnitude of the mutability decline is arbitrary because it depends on the starting sequence, which in this case is random. It also took a large number of mutations to reach this equilibrium (mean 2 mutations per site). This long time to equilibrium likely results from the fact that the non-coding DNA was random and therefore quite far from its equilibrium. In reality, if a given genome is near its exon mutability equilibrium, small changes to the mutation spectrum are unlikely to require evolution to an altogether different equilibrium, and therefore would not take nearly as long. The overall reduction in mutability implies that high mutability trinucleotides were mutated more frequently, and the lower mutability trinucleotides rose in frequency. A general negative correlation between the mutability of trinucleotides and their frequency at equilibrium is evident, especially in highly mutable trinucleotides that contain CpG sites. However, the patterns of trinucleotide change through our simulations are complex, which we discuss more thoroughly below. Our results support that in the absence of other forces, the influence that sequence context has on mutability should lead DNA to settle at an equilibrium that reflects the least mutable sequences.

The premise of our model was that low mutability sequences will become more common because they tend to be retained for a longer time in the genome; however, the observed pattern was more complex. Early in the simulation, the number of times a trinucleotide was chosen for mutation increases with mutability (Figure 1A: black). Counterintuitively, this does not result in a clear negative relationship between the mutability of a given trinucleotide and its change in frequency in the simulation (Figure 1C). There are a few mechanisms that we believe are obscuring this relationship. First, although the rate at which a trinucleotide is removed by mutation depends on its individual mutability, the rate at which a trinucleotide is created depends on the frequency and mutability of the trinucleotides that generate it (Figure 1A: grey). This means that trinucleotide change over time is not a simple relationship with the probability of mutation. Second, when mutations occur, they also alter the mutability of the flanking trinucleotides. This can further obscure the relationship of mutability with equilibrium frequency when, for example, a low mutability trinucleotide is not stable because its flanks have a high mutability (e.g. G<u>A</u>C has mutability within the lowest 1.5% of all trinucleotides, however it decreases in frequency during the simulation, likely due to changes when it’s flanked by the mutable A<u>C</u>G trinucleotide). These processes are further complicated over time as mutations occur on top of each other. This creates an effect where later in the simulation the number of times a triplet is mutated no longer correlates to its mutability (figure 1B:black), the result being that it is not necessarily true that low mutability trinucleotides will be at higher frequency at equilibrium (figure 1D). Similarly, Sung et al. showed that trinucleotide composition correlated with mutation rate in bacteria with small effective populations size. These interactions among sites, even with relatively small windows of sequence context (ie. trinucleotides), highlight the complexity of this process and the importance of using simulations. Despite these intricacies the overall pattern of a lower mutability equilibrium remains clear.

**Figure 1:**
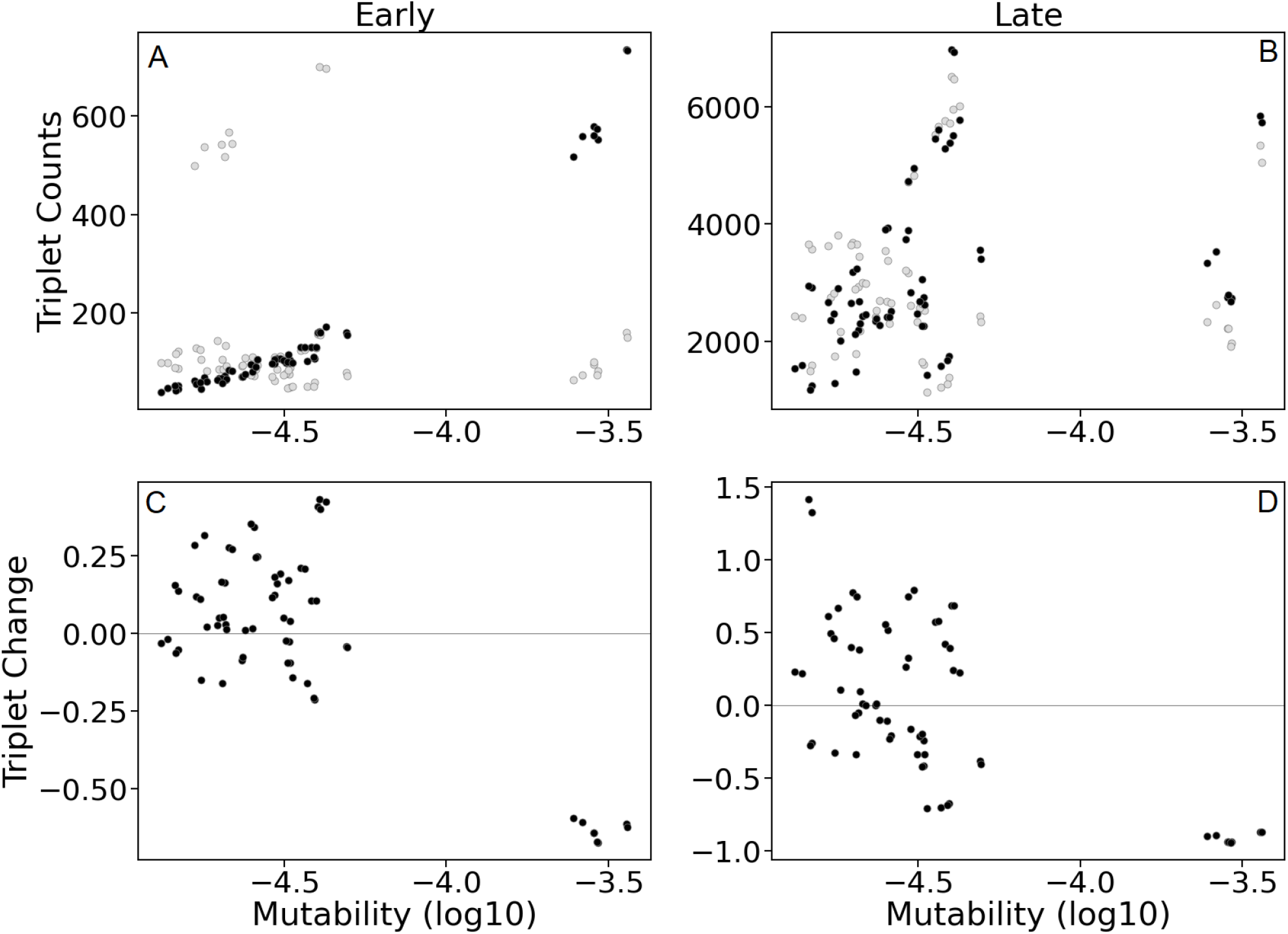
Relationship of trinucleotide mutability with trinucleotide variation in early (10% mutational coverage) and late (200% mutational coverage) stage simulations. Panels A and B display the number of times a trinucleotide is chosen to mutate in the simulation (black) and how many times a trinucleotide is produced from a mutation event (grey) in early (A) and late (B) stage simulations. Panels C and D display average trinucleotide change proportional to initial frequencies in early (C) and late (D) stage simulations. Grey horizontal line denotes the zero mark, indicating no change from initial state. Data for all panes was generated using ~100bkp of randomly generated sequence with no purifying selection (n=10).

### Can purifying selection increase mutation rate?

Our second prediction was that purifying selection in coding sequences stops some mutations from fixing, resulting in the retention of more mutable trinucleotides and a higher equilibrium mutability than neutral sequence. To test this prediction, we inserted random human exons in non-coding DNA to compare mutability change between these two sequence classes. To simulate purifying selection, non-synonymous mutations were accepted or rejected in proportion to the BLOSUM90 matrix. Exon mutability decreased to an equilibrium value 18% higher than non-coding sequence (Figure 2). Our result confirms the prediction that purifying selection generally restricts the ability of sequence to evolve to a lower mutability state. A similar pattern was simulated by Rong et al. when modelling the evolution of splice sites. The fact that coding sequence had higher mutability was not solely driven by the highly mutable CpG trinucleotides as simulations removing CpG trinucleotides produced a qualitatively similar result. Within coding sequence, the effect of purifying selection can be seen at the trinucleotide scale, as mutability has only a weak effect on the change in codon frequency through the simulation (*adjusted R^2^* = 0.16, *p* << 0.01) as compared to the effect of mutability on trinucleotide change within non-coding sequence (*adjusted R^2^*= 0.37, *p* << 0.01). We also tested whether mutability differences among the trinucleotides might interact with purifying selection to create codon usage bias (Jia and Higgs 2008; Powdel et al. 2010), but found no significant relationship between equilibrium codon bias in our simulation and true codon bias in the human genome (*R^2^* <0.001, *p* ~0.5). Note that in the sliding window of mutability presented in figure 2 we observed that some exons had relatively low equilibrium mutability (e.g., first and last exon in figure 2), which appear to be exceptions to our general finding. However, in all these cases the exons started with low mutability and experienced minimal mutability reduction during the simulation (only 33% the amount of decrease experienced by the other codons). Thus, even low-mutability exons demonstrate that coding sequences are limited in their ability to evolve towards a lower mutability state due to purifying selection.

**Figure 2:**
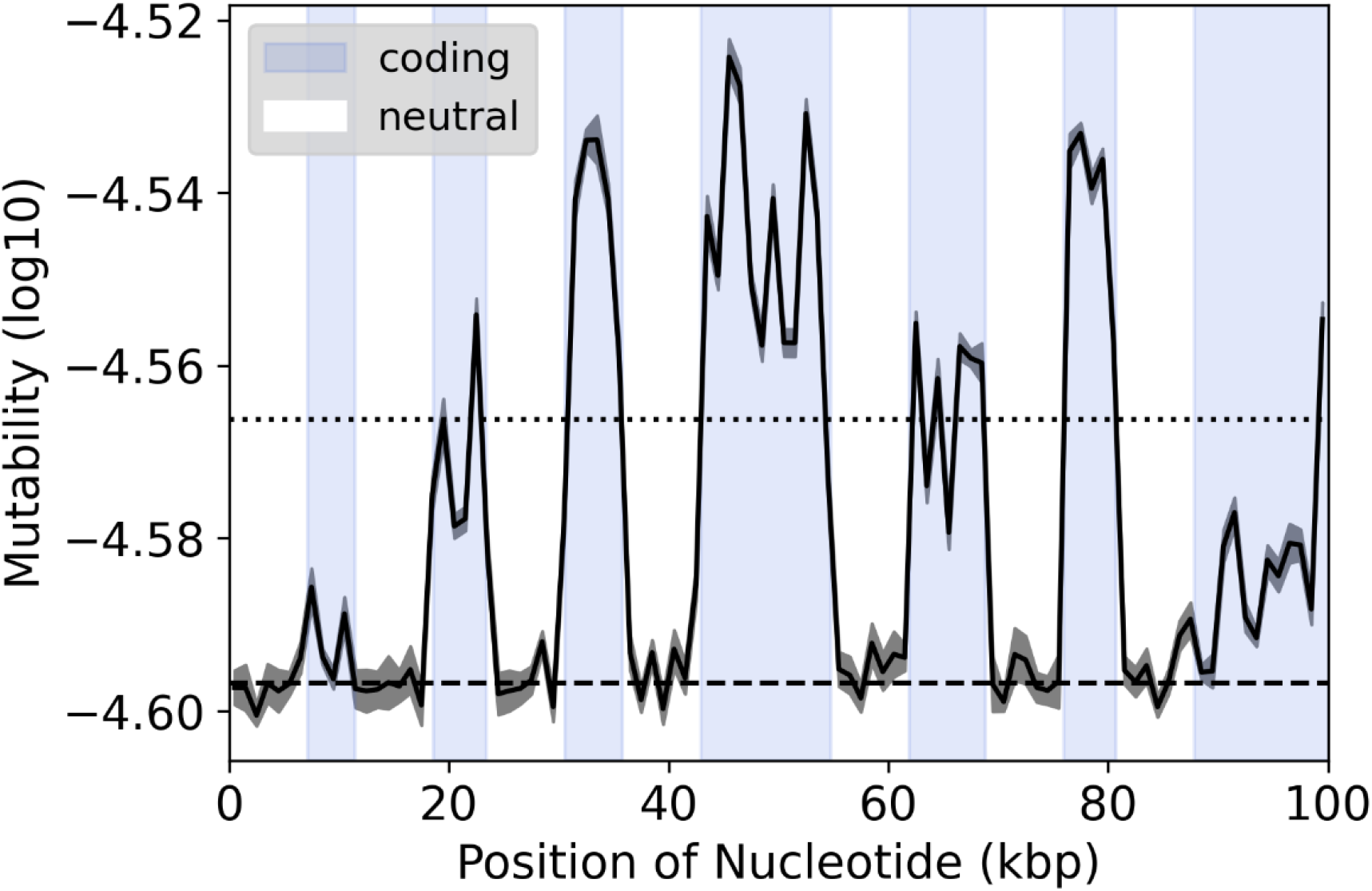
Sliding window of mutability from a simulated chromosome. The chromosome consists of ~100kbp of coding (blue) and non-coding (white) regions at a ~1:1 ratio in an alternating pattern, simulated for 200k iterations. Mutability is calculated as log10 values from the frequencies of trinucleotides in non-overlapping windows of 1kbp. Dark grey ribbon represents standard error between replicate simulations (n=10). The dotted and dashed lines represent the average mutability for all coding and non-coding regions, respectively.

One might expect that coding regions would evolve lower mutability than non-coding regions to reduce the harmful consequences of mutation in functionally important regions. Our simulation shows the opposite, that purifying selection could in fact increase mutability if sequence context is a driver of mutation rate variation. In support of our findings, when we used our mutation model to estimate mutability in intergenic vs. coding regions within the human genome, we found that exons had ~40% higher mutability than intergenic regions. Furthermore, the non-coding mutability reached in our simulations was within 9% of the observed mutability in human intergenic sequence, despite starting from a random assortment of nucleotides. Similarly, at equilibrium, the mutability of coding exons in our simulations was within 22% of true coding sequences in the genome, despite experiencing more complex functional constraints and evolutionary forces not represented in our model. The extent to which true mutation rate variation is explained by the processes simulated here will depend on the relative role that sequence context has on mutability. We know germline mutation rates have been associated with other genomic features such as histone markers, transcription rate and replication timing (Chen et al. 2017; Supek and Lehner 2019). However, Michaelson et al. showed that the effect of trinucleotide sequence on mutability was 5 times greater than the next strongest predictor. Despite the potential to generate more precise models of mutability, the general principle that high mutability sequence will tend to destroy itself means that tension between mutation and purifying selection could prove to be fundamentally important to our models of how mutation rate variation evolves.

## Methods

One of the best predictors of mutation rate at a single site is the base itself and the 2 bases immediately flanking it (trinucleotide). Here, we used 108k single base mutations from the human germline (Jónsson et al. 2017) and their flanking bases to estimate a mutability model of the relative mutation rate (mutability) at each trinucleotide as well as the probability of the focal site mutating into each of the three other possible nucleotides. Trinucleotide mutability was calculated as the number of mutations of a given trinucleotide over the number of those trinucleotides present in the human reference genome (Hg38). For each trinucleotide we also assigned the probability of each of the three possible mutations based on the frequency of each change in the observed mutation data.

To simulate sequence and mutability evolution over time, we started with a hypothetical chromosome and mutated it according to the mutability model. The initial chromosome consisted of ~100kbp of coding and non-coding regions at a ~1:1 ratio in an alternating pattern. Non-coding sequence was randomly generated from equal proportions of the four nucleotides. To represent true coding sequence and the higher-order patterns of codon and amino acid usage we used random human coding sequence (CDS) (version Hg38). To represent purifying selection on amino acid sequence we transformed the BLOSUM90 matrix such that the probability of a mutation fixing was proportional to its score. Additionally, to represent the presence of sequence critical to protein function we randomly assigned 50% of the coding sequence as invariant so that no mutations could occur. Each iteration of the simulation followed the following steps:

1. Assign mutability of each position in the chromosome based on its trinucleotide sequence.
2. Randomly sample one position weighted on all the mutability of all sites.
3. For the chosen base, assign which of the three possible mutations occurred with weighted probabilities.
4. Accept or reject the proposed mutation using the following criteria:

a. If the mutation is in a non-coding region, accept the change.
b. If the change is in an invariant site, reject the change.
c. If the site is in a coding region but not an invariant site, then the proposed mutation is accepted or rejected with a random probability that is proportional to the score from a BLOSUM90 matrix.

These steps were repeated, recalculating mutability after each iteration, until mean mutability reached equilibrium. The number of iterations was set to 200k cycles, the point at which we determined the sequence enters equilibrium (no large changes in mutability, see figure S1). To further investigate particular non-coding mutational patterns, we also ran separate simulations with a 100Kbp sequence consisting of only non-coding DNA.

We also explored the influence that sequence-dependent mutability would have on the relative frequencies of synonymous codons, or codon usage bias. To investigate this hypothesis we calculated codon usage bias in the genome and used modified human CDS to understand whether sequence context mutability alone plays a role in creating codon usage bias. To this end, we used ~100kbp of cumulative sequence from randomly selected exons 4-6kbp in length. For each amino acid in the sequence we replaced the existing codon with a random codon from the sample of synonymous codons for that amino acid; the result being a sequence without codon usage bias that still codes for the same amino acid. This sequence was mutated for 200k cycles according to the rules outlined in steps 1 through 4 above.

## Notes

### Competing Interest Statement

The authors have declared no competing interest.

