## Supplemental figures for "How sequence context-dependent mutability drives mutation rate variation in the genome"

### Supplemental information:

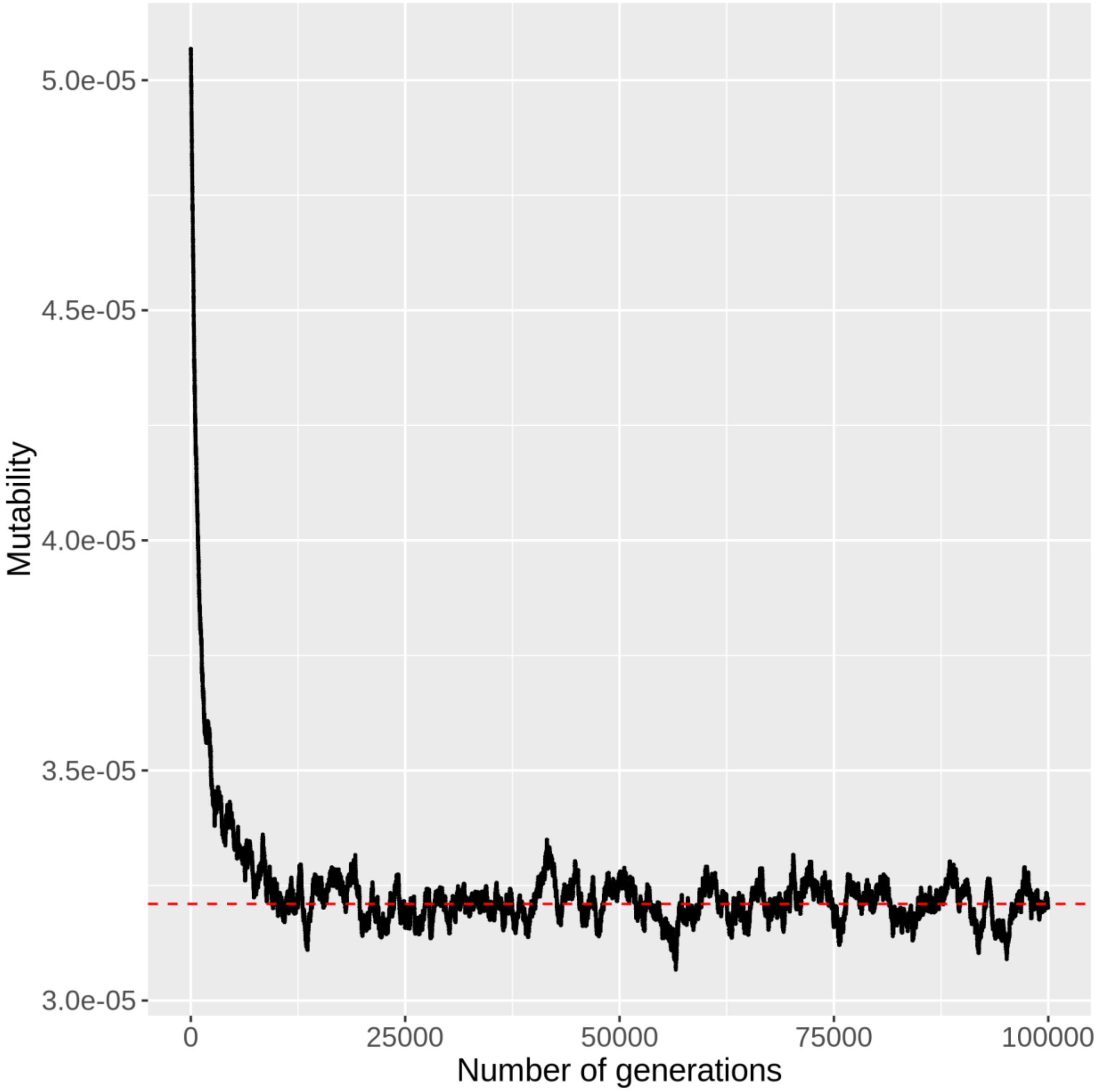


Figure S1: Line plot of mutability's exponential decay over time in a 10kbp chromosome. The chromosome consists of coding and non-coding regions at a ~1:1 ratio in an alternating pattern, simulated for 100k generations. Global mutability reaches an approximate equilibrium point of 3.21e-05 after 20k generations (200% mutational coverage).

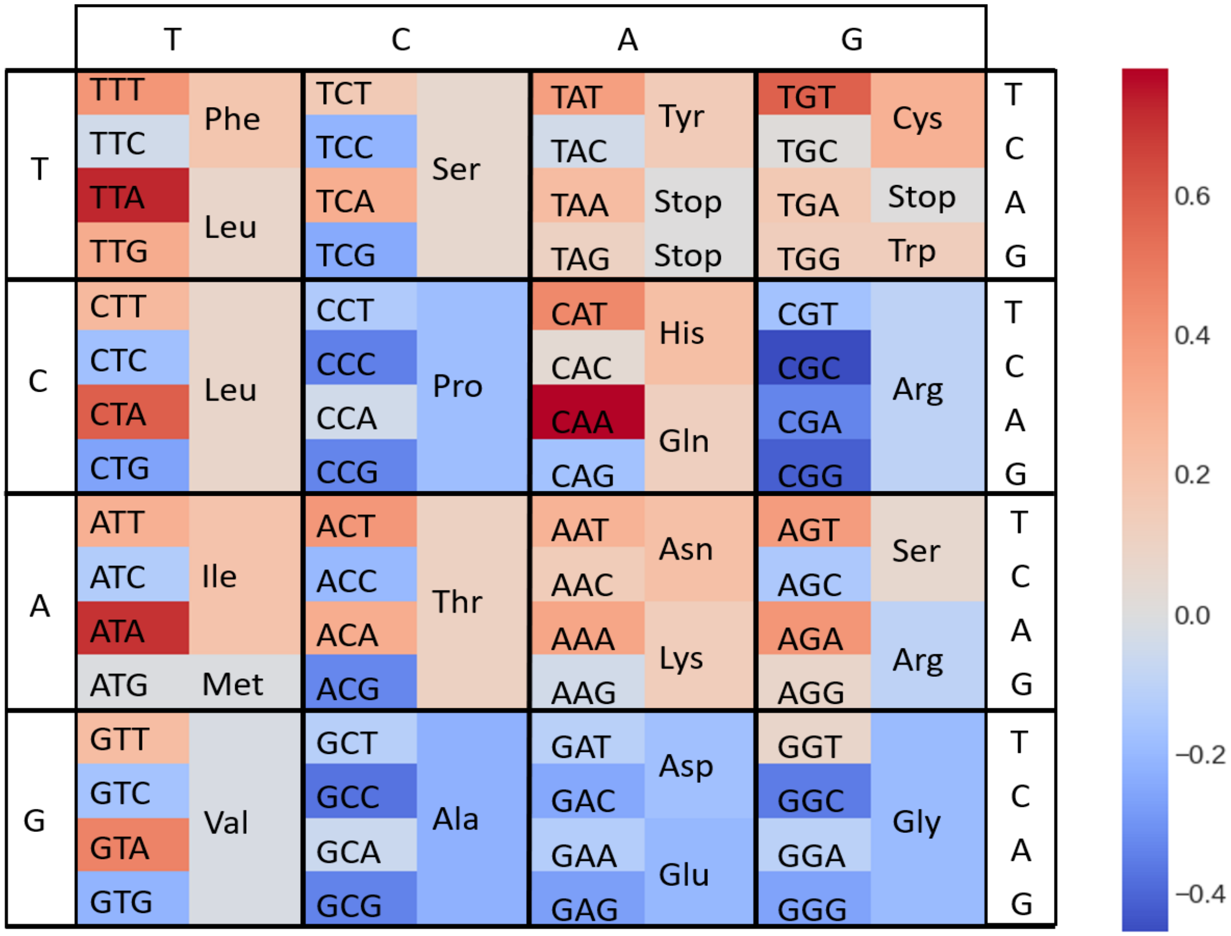


Figure S2: Heatmap of average codon and amino acid change proportional to initial frequencies in the coding sequence. Average change was calculated from 10 trials of different simulated chromosomes with 200% mutational coverage, each with ~50kbp of coding sequence from 8-9 randomly chosen human CDS.
